# Counter-intuitive method improves yields of isotopically-labeled proteins expressed in flask-cultured *Escherichia coli*

**DOI:** 10.1101/2024.05.15.594275

**Authors:** Miguel Ángel Treviño

## Abstract

NMR is a powerful tool for the structural and dynamic study of proteins. One of the necessary conditions for the study of these proteins is their isotopic labeling with ^13^C, ^15^N and sometimes ^2^H. One of the most widely used methods to obtain these labeled proteins is heterologous expression of the proteins in *E. coli* using ^13^C-D-glucose and ^15^NH_4_Cl as the sole nutrient source. In recent years, the price of ^13^C-D-glucose has almost tripled, making it essential to develop labelling methods that are as cost effective as possible. In this article, different parameters have been studied to achieve the most rational use of ^13^C-D-glucose and an optimised method has been developed to obtain labeled proteins with high labelling, low ^13^C-D-glucose consumption. Surprisingly, the optimised method is also simplier and does not require monitoring culture growth

## INTRODUCTION

An essential step to study proteins by NMR is to obtain ^13^C, ^15^N and/or ^2^H labeled proteins at high concentrations to transmit magnetization between the different nuclei during the NMR experiments.

The most widely extended method to obtain high quantities of proteins for biological studies is to produce them in heterologous systems, mainly in *E. coli*, a very well-studied system with many variants and strains. Although biofermentors allow fine, real-time control of many of the variables affecting the growth of the bacteria and the protein expression, they are expensive and the majority of laboratories use simpler types of equipment such as Erlenmeyers or other flasks shaken in incubators. Using these low-tech equipments, enough protein can be obtained in most cases. For unlabeled proteins, the cells are grown in a rich medium (LB, 2YT, or other) until the cells are growing at an exponential rate -usually at optical density between 0.6 and 0.8 units-. Then the appropriate inducer is added to the medium to initiate the expression of the selected protein.

In the case of labeled proteins, instead, the classical protocol uses a modification of M9 minimal medum (Miller 1972) containing ^13^C-D-glucose and ^15^NH_4_Cl as solely carbon and nitrogen sources, respectively, instead of the rich medium, maintaining the point of induction at 0.6-0.8 OD600.

Over the years, many modifications have been incorporated into this basic protocol trying to improve the yield of labeled protein due to the high cost of the isotopic sources. So, two main strategies have been followed: 1) modifications in the minimal medium and growth conditions 2) previous generation of non-labeled biomass The first strategy is focused on controling variables such as medium pH which usually decreases during bacterial culture. At acidic pH, the cells stop growing and enter in stationary phase and if pH is lower than 4.5 can even prevent further cellular growth (Neidhardt et al. 1974; Sánchez-Clemente et al. 2018). To prevent this effect, the concentrations of buffering substances can be increased to the limit of solubility ((Neidhardt et al. 1974)(Cai et al. 2019)). To keep the cells in exponential growth phase, it is necessary to maintain the bacteria in aerobial conditions, usually achieved with a high rate of agitations or using baffled recipients to generate a turbulent flow of the medium instead of a laminar one. Recently Cai et al(Cai et al. 2019) have published a method in which they reduce the culture temperature. This would increase the dissolved oxygen by around 10% when the culture is kept at 30ºC or around 20% at 25ºC, compared to the usual 37ºC water solubility(Bok et al. 2023). In addition, lower growth rates at cooler tempertures increases the number of ribosomes per cell, raising the proportion of ribosomes available to produce the protein of interest(Marr, 1991).

Regarding the second strategy, in the seminal paper from Marley et al (Marley et al. 2001), they grew bacteria up to 0.6 OD_600_ and then centrifuged and changed the medium to a minimal one, diminishing the volume to reduce isotope consumption. After 1h of adaptation for incorporation of the isotopes and generation of labeled amino acids, the protein expression is induced. Variations over this protocol have been proposed. For example, Sivashanmugam *et al* (Sivashanmugam et al. 2009) grew bacteria up to 3-5 or 5-7 OD_600_ in rich media before centrifugation and swapping to minimal medium (in this case without volume reduction). Similarly to Marley et al, cells are kept for 1-2 hours in minimal medium before induction to ensure the incorporation of the isotopes into the precursors of the protein.

In both strategies a fraction of the isotopes are consumed for generation of biomass (strategy 1) or to ensure complete isotope incorporation and adaptation to the minimal medium (strategy 2), and it is not harnessed for labeled protein generation.

In this paper, a method that eliminates the necessity of OD_600_ monitoring and centrifugation is presented. This eliminates a stressful step for the bacteria. Also, the conditions to minimize the non-productive consumption of isotopes have been studied, increasing the yield of protein without sacrificing the isotopic incorporation, which is maintained around 98% for ^13^C.

## MATERIALS and METHODS

### Plasmids and E. coli strain

To analyze each variable model *E. coli* BL21(DE3) bacteria transformed with a pET24 plasmid containing the codifying sequence for the human CB1 Cannabinoid Receptor Interacting Protein 1 (CNRIP1) (uniprot Q96F85-1) and containing an Nt-histidine tail and a cleavage site for TEV protease was used.

### Culture conditions

Transformed bacteria were grown in a slightly modified M9++ medium (Cai et al) (table 1). in Erlenmeyer’s or Tunair (IBI Scientific, Dubuque, IA) flasks with capacity at least 10 times bigger than the medium volume. To generate inocula, the transformed strain was grown in LB overnight at 37ºC. The grown inoculum was directly added at the minimal medium.

**Table 1.** composition of modified M9++ minimal growth medium.

| **Table 1. composition of modified M9++ minimal growth medium** |  |
| --- | --- |
| Component | concentration |
| LB medium | 1/1000 |
| Trace elements | 1/5000 |
| MgSO<sub>4</sub> | 1mM |
| 100xBME vitamins solution | 1/400 |
| Thiamine | 10µg/ml |
| CaCl<sub>2</sub> | 0.2mM |
| K<sub>2</sub>HPO<sub>4</sub> | 109.1mM |
| KH<sub>2</sub>PO<sub>4</sub> | 36.7mM |
| Na<sub>2</sub>HPO<sub>4</sub> | 13.8mM |
| D-glucose or <sup>13</sup>C-D-glucose | Variable<sup>1</sup> |
| <sup>15</sup>NH<sub>4</sub>Cl | Variable<sup>1</sup> |

<sup>1</sup> Different percentages of nutrients were tested. Nutrients needed for the final protocol after optimization are: D-glucose 0.2%; <sup>15</sup>NH<sub>4</sub>Cl 0.06%. After biomass generation, additional <sup>13</sup>C-D-glucose 1% and <sup>15</sup>NH<sub>4</sub>Cl 0.3% are added.

The cultures were composed of three phases: 1) biomass generation in minimal media with non-labeled glucose, 2) addition of extra glucose (labeled or unlabeled, depending on the tested conditions) for biosynthesis of amino acids and culture for incorporation of isotopes, and 3) induction and protein expression.

The temperatures and times in each step were 25ºC overnight (phase 1), 30ºC a variable time (phase 2) and 20ºC 24 hours (phase 3), except when indicated.

Cultures for other methods, to comparisons, were performed as described in their original papers (Cai et al (2019), Sivashanmugam et al (2009), Marley et al (2001))

### Culture Variable determinations

Optical density was measured at 600nm in a Nanodrop One spectrophotometer (Thermo Scientific, Waltham, MA).

Free D-glucose concentration in the media was determined using a commercial QuantiChrom Glucose Assay Kit (Bioassay Systems, Hayward, CA), which generates a blue color due to the formation of an imino bond between the aldehyde group for sugars and o-toluidine. After a centrifugation pulse of the culture to eliminate the bacteria, 2 to 5 microliters of the supernatant were tested with 50ul of reactive, following the manufacturer’s instructions. Measures for A630 were performed in a Nanodrop One spectrophotometer and D-glucose concentration was calculated by interpolating in calibration curves obtained during the same experiment.

### Expressed protein quantification

Volumes corresponding to equivalent amounts of D-glucose added during phase 2 (i.e., 100μL for 1% glucose added, 200μL for 0.5% glucose added or 250μL for 0.4% glucose) of final expression cultures were centrifuged and pellets were lysed with 50 μL bugbuster (Merck, Rahway, NJ). After centrifugation, the insoluble fractions were solubilized in 50 μL 8M urea. Both fractions were pooled and mixed with the same volume of loading buffer for PAGE. 2 to 10 μL were loaded in homemade gradient (4-25%) acrylamide gel containing 3.75% trichloroethanol for direct fluorescence detection of triptophans (Kazmin et al. 2002) and the proteins were separated by PAGE. Each sample was loaded at least 3 times.

Bands were quantified in a ChemiDoc MP Imaging System (Biorad, Hercules, CA) using the free stain option and analyzed using the ImageLab software.

### Protein purification

For mass spectroscopy experiments, bacteria from 5ml growths were centrifuged and resuspended in 1 mL of potassium phosphate 50mM pH8, NaCl 300mM, and imidazole 10mM with 1 μL of Halt inhibitors (Thermo Scientific, Waltham, MA). Cells were sonicated and the lysate was centrifuged. One hundred μL of Nickel High Density beads (Agarose Bead Technologies, Torrejón de Ardoz, Spain) were added and the mixture was loaded in a MicroBiospin empty column (Biorad, Hercules, CA). After washing with the same buffer, CNRIP1 was eluted in 400 μL of potassium phosphate 50mM pH8, NaCl 300 mM, and imidazole 500mM. Two μg of TEV protease were added and the sample was dialysed against 1L of potassium hydrogen phosphate 5mM p? 6.8, NaCl 10mM, β-mercaptoethanol 1mM.

For NMR spectroscopy, the cultures were scaled up to 50 or 100mL. Lysis was performed analogously but the lysate supernatants were loaded in HisTrap 5mL FF columns (Cytiva,,) the eluates were dialyzed in potassium phosphate 5mM pH8, NaCl 10mM and simultaneously cleavaged with TEV protease. The dialyzed samples were loaded in the same column at the flowthrough collected, and re-dyalized. Samples were then loaded in HiTrap 1mL SP columns. The samples were prepared in 5mM potassium phosphate pH 6.8, NaCl 10mM.

### Mass spectrometry

The mass of the purified proteins was determined by mass spectrometry. Samples were analyzed in a HPLC 1100 Series LC System (Agilent Technologies, Palo Alto, USA) coupled to an HTC-Ultra ETD II ion trap mass spectrometer (Bruker Daltonics, Fremont, USA) using an electrospray ionization (ESI) source. The molecular masses of proteins were calculated by deconvolution of the ESI-MS spectra using the Thermo Finningan BIOMASSTM software (Thermo Fisher Scientific, San José, CA, USA).

### NMR spectroscopy

1D ^1^H-spectra and ^1^H-^13^C-HSQC spectra were recorded on a Bruker Avance Neo 800 MHz (^1^H) spectrometer fitted with a cryoprobe and z-gradients. The experiments were collected at 25°C.

For coupled spectra, the same experiments were performed as for conventional decoupled spectra, but no ^13^C decoupling pulses were applied during acquisition.

## RESULTS and DISCUSSION

Previous work (Cai et al. 2019)has shown that in *E. coli* cultures at low temperatures, up to OD_600_=6, before induction, a high amount of protein is obtained (relative to the amount of medium used), with isotopic labeling of about 97%. Despite this good result, there is a percentage of the ^13^C-D-glucose that is used just to generate biomass and therefore it is “wasted” to improve the yield of labeled protein.

A simple way to improve this would be to combine this protocol with a first step of biomass generation in rich media, just like other protocols, which then switch to labeled media by centrifugation and produce high yields of labeled proteins, but centrifugation steps can be stressful for the bacteria and they recover slowly. In fact, the OD_600_ can drop in the first few moments in minimal media and it is difficult to estimate how long the bacteria need to be grown in this media before induction to maximise expression and minimise detrimental unproductive consumption of labeled nutrients.

Therefore, it can be hypothesised that complete consumption of unlabeled glucose in minimal media could be as efficient in terms of biomass production as using rich media but avoiding the stress of centrifugation. It has been reported that *E. coli* recover quickly from short periods of starvation with no apparent sequelae (Lempp et al. 2019). In addition, the cells would be adapted to grow in these minimal media from the start, further reducing the stress of switching from rich to poor media and minimising the time required to incorporate labeled metabolites.

To evaluate this hypothesis, biomass production and glucose consumption were monitored in different conditions (**Figure 1**). Different initial D-glucose concentrations were tested. In all cases, after 23h growth at 25ºC, 0.5% D-glucose was added and continued to grow at 30ºC for 1.5h followed by growing at 20ºC for other 23h. The depletion of glucose after overnight growth was complete in all conditions tested and growth rate recovery appeared to be complete after the supplemental 0.5% D-glucose addition. Although it was predictable that some of this additional D-glucose would be consumed during the isotope integration step, after 1.5h at 30ºC the remaining nutrient is extremely low for the cultures with high initial glucose concentrations, leaving less than 0.1% D-glucose available for the protein expression step in the condition of initial 0.5% D-glucose and, after 3 additional hours at 20ºC, there is not remaining glucose for any of the initial glucose conditions tested.

**Figure 1:**
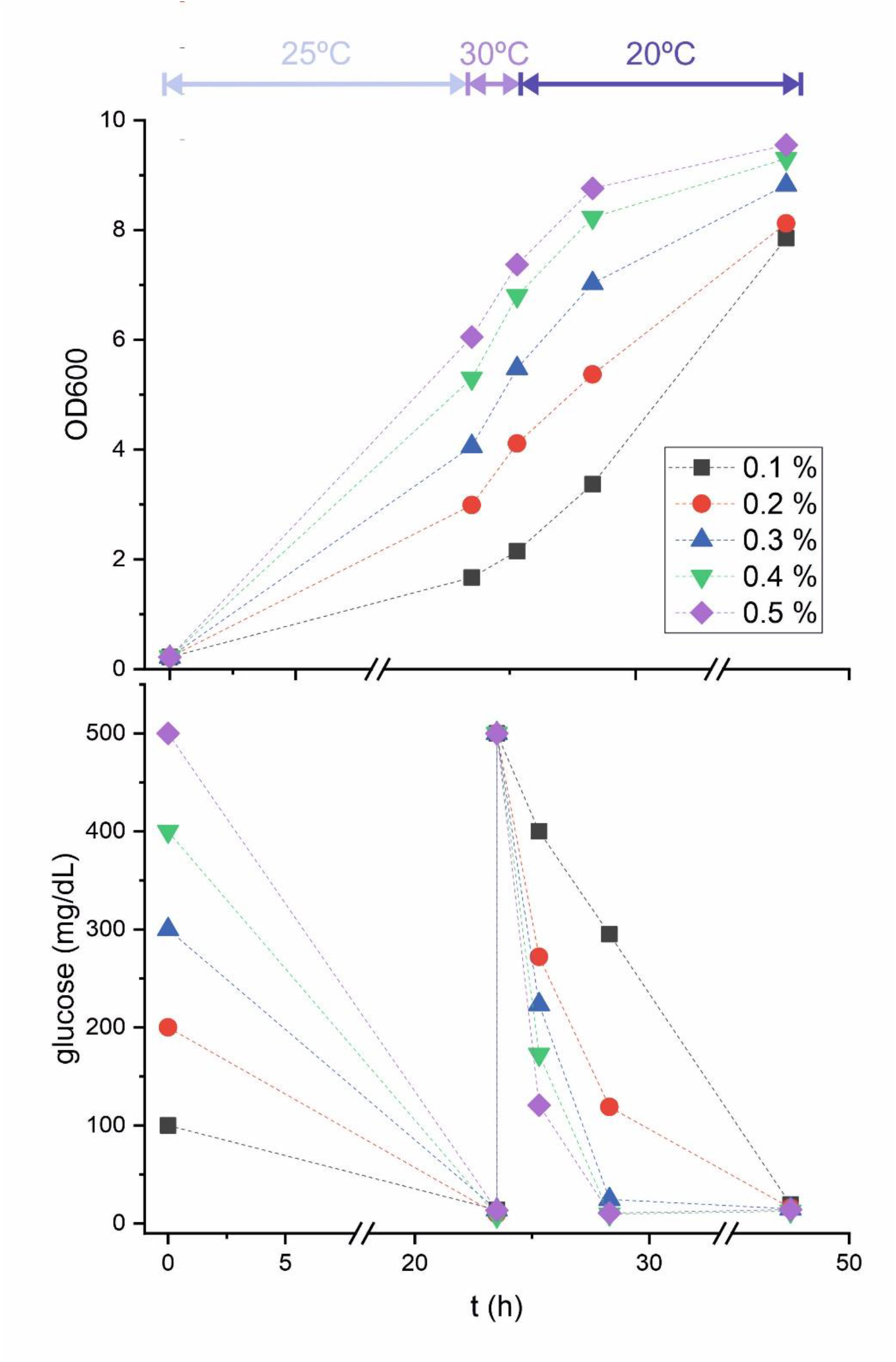
Optical density (top) and dissolved D-glucose (bottom) in *E. coli* cultures as a function of initial D-glucose concentration. Cultures were kept at the temperatures indicated at the top of the figure, mimicking the steps in the induced cultures, i.e. 25ºC during the ‘biomass generation step’, 30ºC after addition of 0.5% D-glucose to simulate the ‘isotope integration step’ and 20ºC to simulate growth after addition of IPTG in the ‘expression step’. Dashed lines do not represent linear growth and are only added to help locate points from the same conditions.

Thus, a counterintuitive result was found: it is not convenient to produce large amounts of biomass, but to find a compromise between biomass and consumption of labeled glucose to improve the final protein yield.

The second variable monitored was the incorporation of ^13^C into the labeled protein. The data show that for high initial glucose only partial incorporation into the protein was reached but when the initial glucose was reduced from 0.5% to 0.4% and to 0.2% incorporation increased from 75% to 90% and 98%, respectively, as detected by NMR (**Figure 2**).

**Figure 2:**
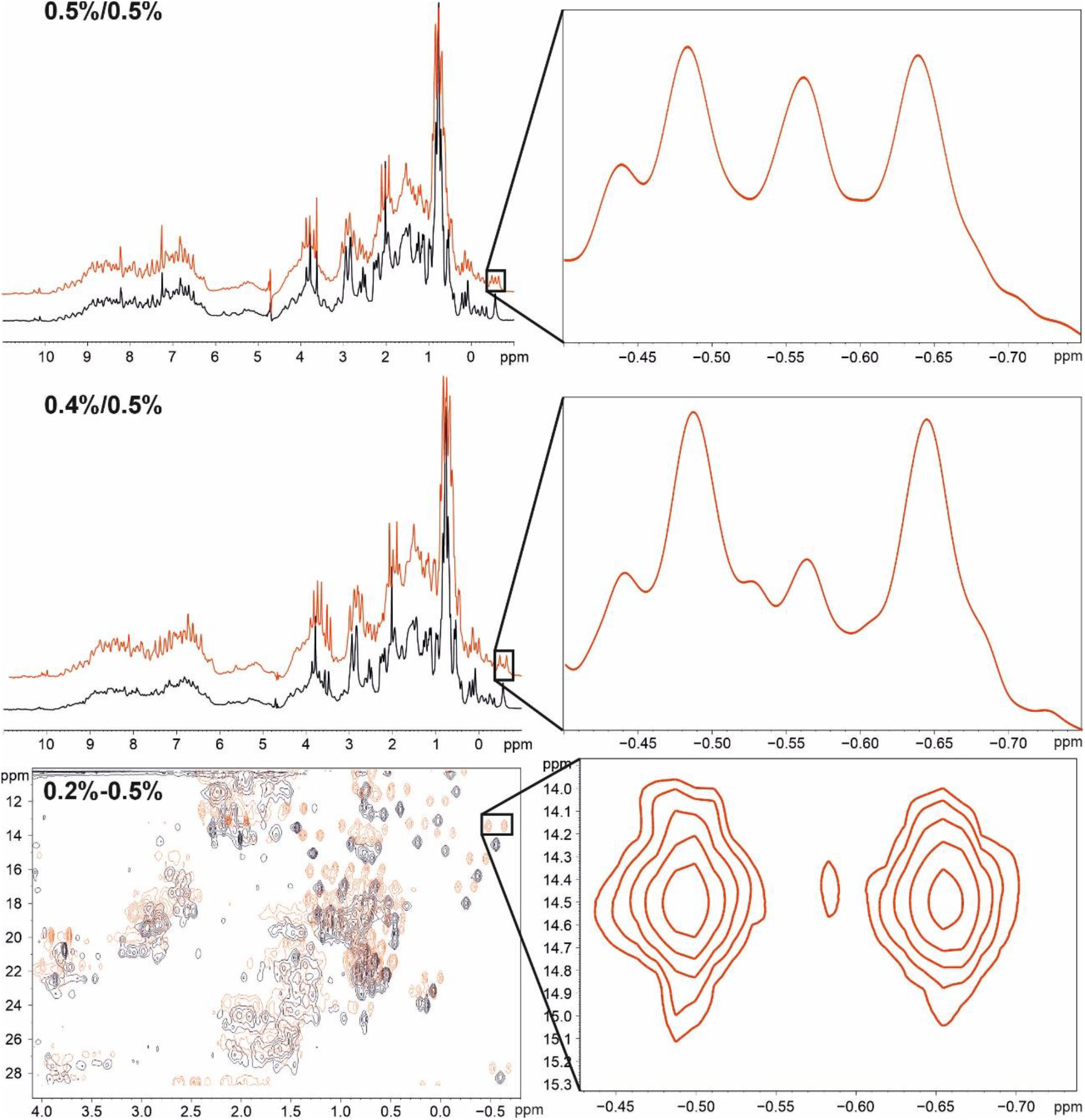
^13^C incorporation at different culture conditions measured by NMR. 1D ^1^H spectra with (black) or without (red) a ^13^C decoupling pulse. On the left, the full spectra are shown; on the right, the selected region is zoomed in. To improve clarity, the coupled spectra (red) are shifted vertically. For conditions 0.5%-0.5% and 0.4%-0.5% 1D NMR spectra are shown. For 0.2%-0.5%, a 2D ^1^H^13^C-HSQC spectrum is shown because its sensitivity and lack of superposition allow better quantification of the signals.

These results led to the testing of other conditions - decreasing the glucose in the biomass generation step and increasing the labeled one added in the “isotope integration step” -. The protein yield in each new condition showed a better yield for 0.2-0.3% D-glucose in the “biomass generation step” combined with 1% D-glucose added in the “isotope integration step” (**Figure 3**).

**Figure 3:**
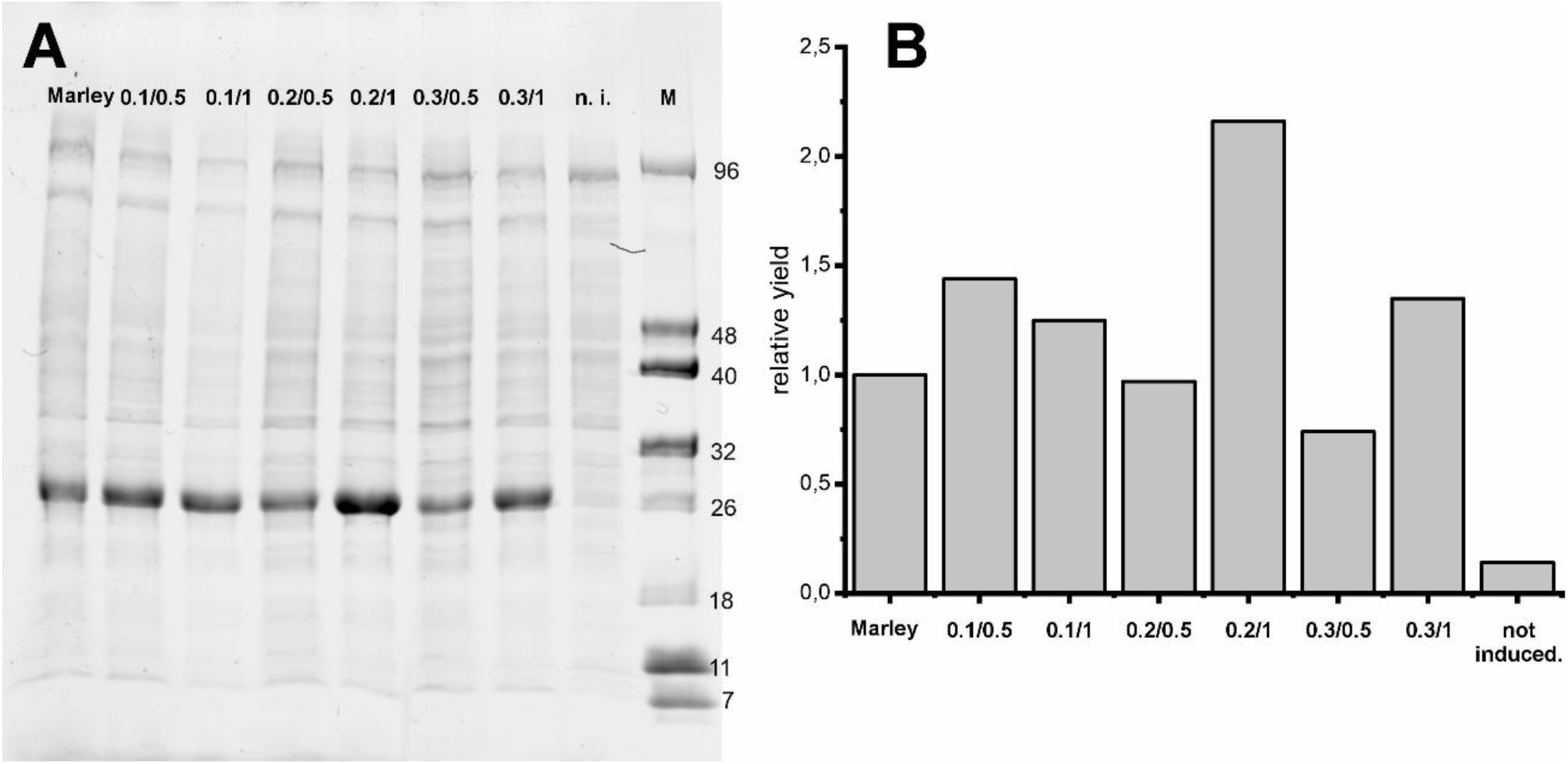
Yield of protein at different percentages of initial D-glucose and added D-glucose. **A**.: Fluorescence emission under UV exposure of SDS-PAGE. **B**.: Bar graph representing the relative fluorescence according to the method of Marley et al. considered as unit.

Considering the data from the previous experiments, the effect of varying the time of the “isotope integration step” was tested by monitoring the protein yield and the ^13^C incorporation (**Figure 4**). A second counter-intuitive fact appeared: there is minimal effect of the length of this phase on the ^13^C incorporation, which is always around 97-98% and can even be eliminated.

**Figure 4:**
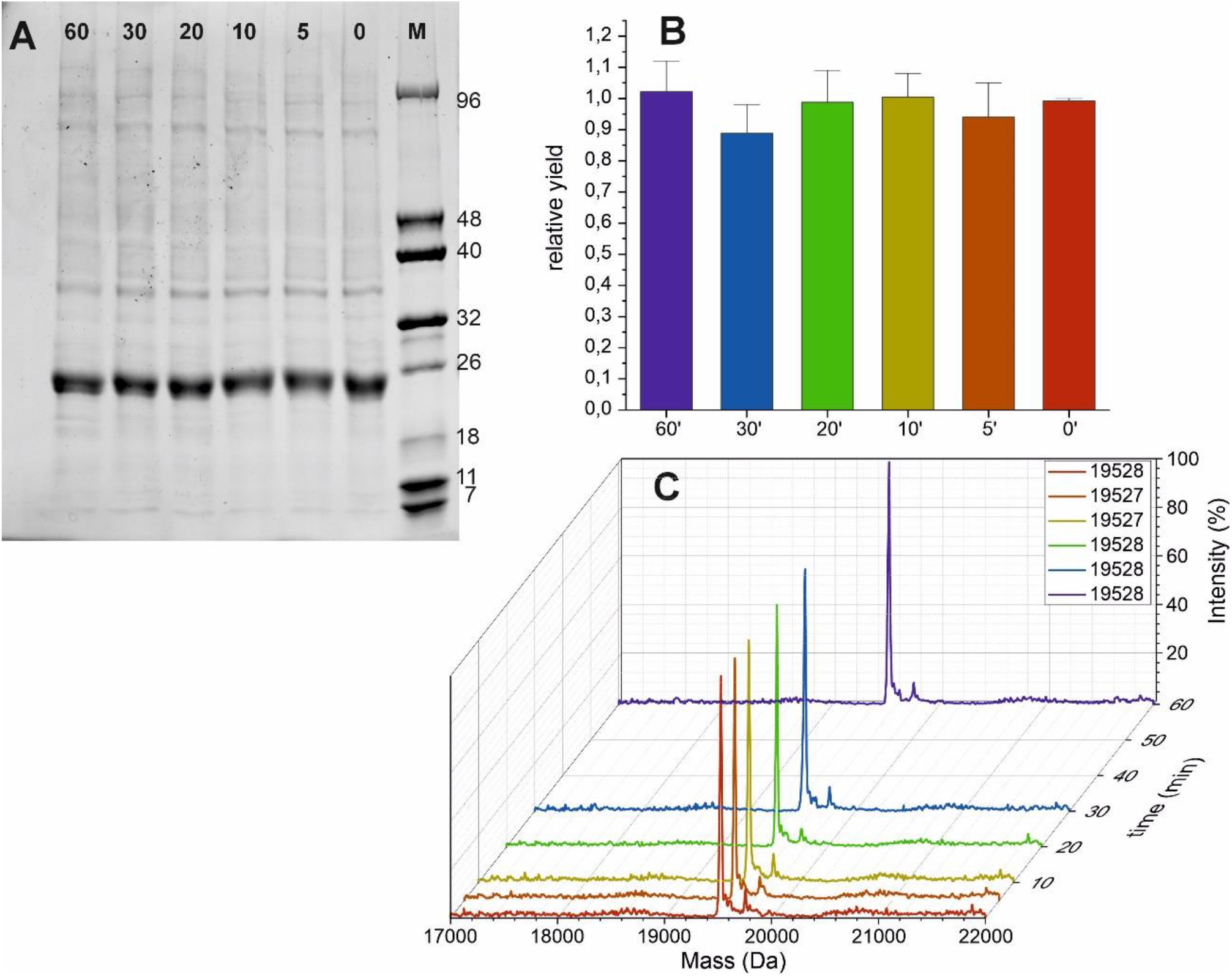
Effect of different “isotope integration” times on yield and ^13^C labelling. **A**: Fluorescence emission under UV exposure of SDS-PAGE. **B**.: Bar graph representing the relative fluorescence according to the method of Marley et al. 2009, which was used as unit. Data are the media of three experiments **C**: Mass spectra. The masses for each condition are given in the table.

As the initial data indicated that for 0.5/0.5% conditions, even with a 1.5h isotope integration step, incorporation was around 75%, It seems that in any case the relative ratio between initial unlabeled and subsequently added labeled glucose should not exceed 20% (0.2% initial D-glucose, 1% labeled D-glucose).

Although all biomass generation cultures were performed at 25ºC to ensure maximum O_2_ available to the cells, the influence of temperature on the growth was monitored at 25ºC, 30ºC and 37ºC (**Figure 5**). As complete glucose depletion is needed for this method, at 25 and 30ºC, overnight culture at 25 ºC or 30ºC is recommended, while at 37ºC complete glucose depletion is achieved after approximately 5.5 hours. This would allow the biomass generation step to be shortened. However, the high growth rate at 37ºC could have some disadvantages, such as a lower number of ribosomes per cell (Marr 1991), or the possible micro-aerobic state, which could promote the accumulation of acetate (Partridge et al. 2007) and therefore inhibit the cellular growth or the expression of the desired protein (Shiloach and Fass 2005). Finally, other *E. coli* strains or plasmid-strain combinations may be less efficient in nutrient consumption than those tested here and may require longer culture times. Although the total time for 25ºC or 30ºC cultures is longer, the active time for the present method is, in fact, decreased lowering from around 1-2 hours (due to measures of OD_600_ until reachs 0.6, centrifugation, resuspension in minimal medium, and addition of inductor) to around 5-10 minutes (due to addition of labeled nutrients and inductor simultaneously).

**Figure 5:**
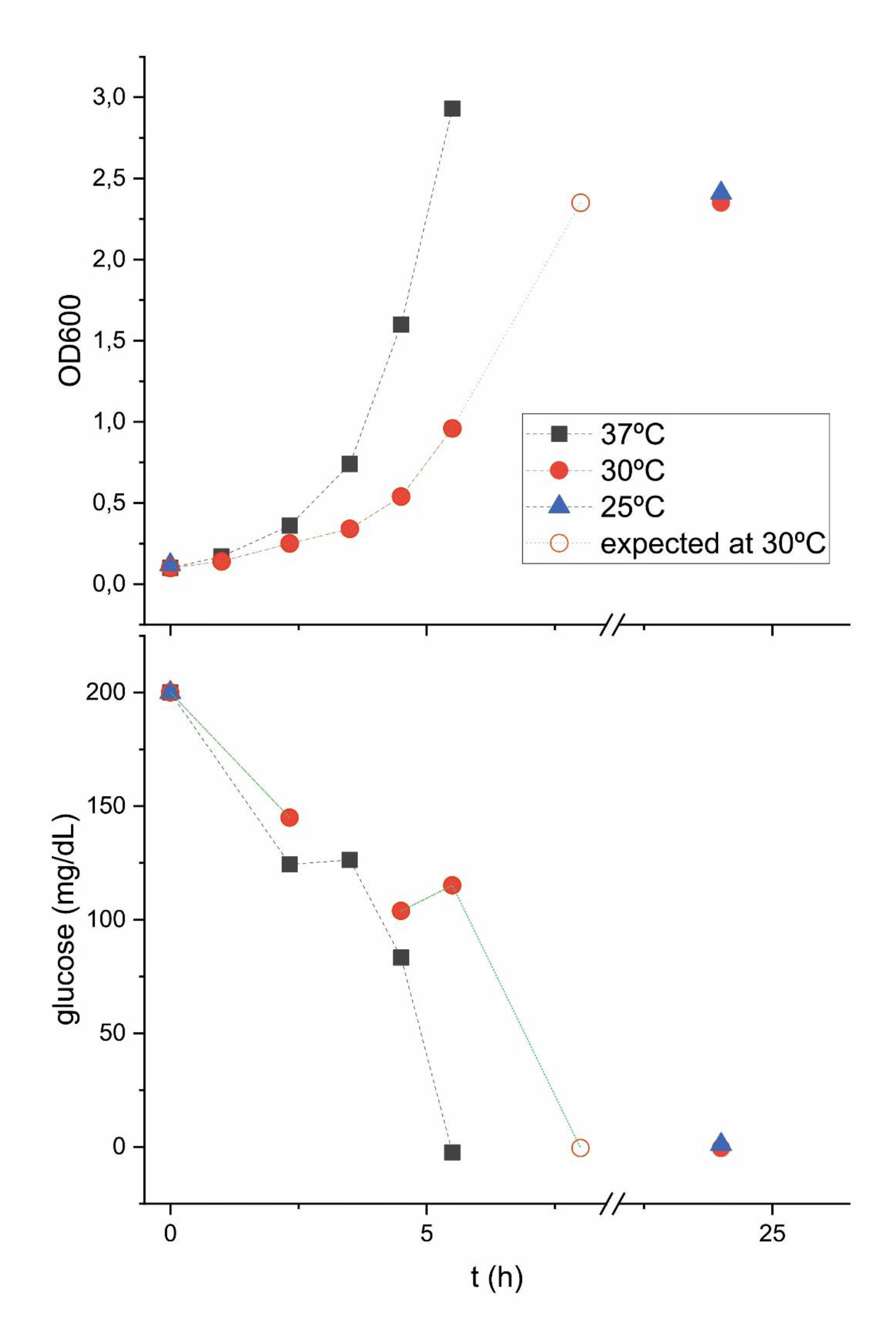
Optical density (top) and dissolved D-glucose (bottom) in *E. coli* cultures as a function of temperature. The dashed lines do not represent linear growth and are only added to make it easier to locate symbols from the same conditions. For 30ºC, no recording was made at the likely hour of complete D-glucose consumption, so an unfilled circle has been added based on the exponential behavior of the curves to give an indication of the approximate time of the event.

The influence on yield of different glucose concentrations during the expression step was also investigated (**Figure 6**). A slight increment of yield seems to be detected when glucose was increased from 1% to 1.5% but decreased at higher percentages. In any case, the increases are in the range of error, so there is not an evident advantage to use higher glucose percentages. In any case, this method clearly improves the yield of labeled protein when compared with some of the previously described protocols and, even in the least favourable case, there is a 20% increase. The tests indicate that a 50 mL culture (0.5g 13C-D-glucose) is enough to obtain 10-20mg of purified labeled protein Although we have not tested the possibility of start the culture directly from colonies from a plate instead of an overnight liquid preinoculum culture, as previously reported (Sivashanmugam et al. 2009), It could also diminish the total time of culture in addition to the already diminished active time.

**Figure 6:**
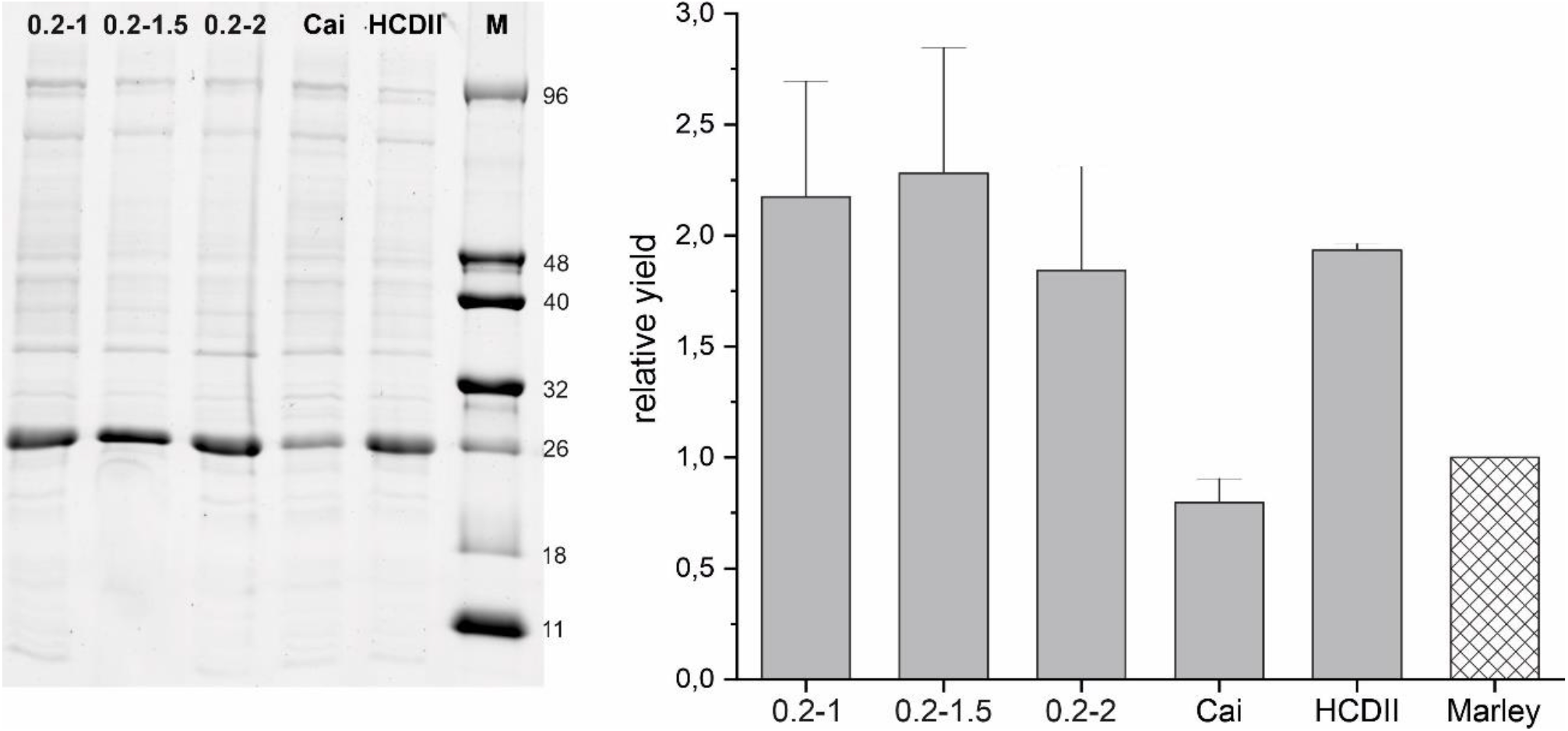
Effect of percentage of D-glucose added in the “protein expression step” on protein yield. A: Fluorescence emission under UV exposure of SDS-PAGE. B.: Bar graph of relative fluorescence. Band intensities were referenced to the HCDII (Sivashanmugam et al (2009)) band. For comparison, a checkered bar corresponding to the relative yield obtained by Marley et al method, determined in previous experiments, was added to the graph. Data are from five experiments.

The ratios of NH4Cl and D-glucose in this recipe are calculated to ensure that glucose is the limiting nutrient. In this way, it can be ensured that all the glucose has been used up in the biomass production step. Since 13C-labeled proteins for NMR are usually also 15N-labeled, 15NH4Cl must to be used from the beginning because it cannot be assured that there is no remaining nitrogen that has not been consumed in the initial steps. Since 15NH4Cl is much cheaper than 13C-D-glucose, net savings still prevail and it will be even more significant if we need to used D7-13C-D-glucose or others isotopic labeled precursors, even more expensive than 13C-D-glucose

## CONCLUSIONS

A new method for the production of ^13^C-labeled protein has been developed (detailed protocol in supplemental material) which minimises the time commitment required (**figure 7**) and optimizes labeled nutrients consumption.

**Figure 7:**
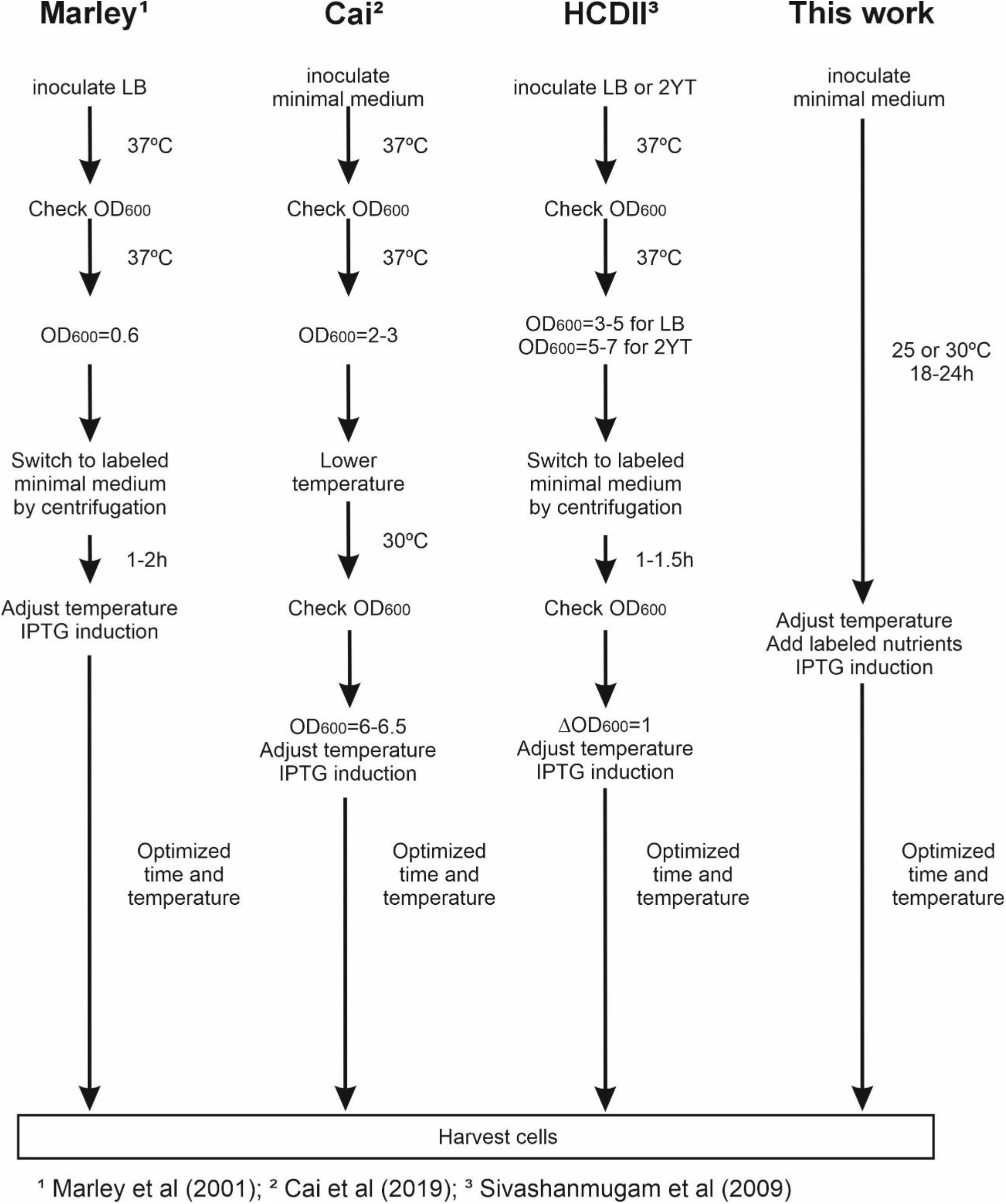
Outline of some protein expression methods compared with the one described here. Different OD_600_ measurements may be required for these methods, increasing the required active time.

An initial culture step at 25-30ºC in minimal medium allows biomass to be generated but avoids centrifugation. No “isotope integration step” is necessary and simply lowering the temperature to the appropriate level for protein expression, before adding labeled nutrients and the inductor, is enough to obtain maximum protein yields with high 13C incorporation This protocol, with minimal variations, could be used to generate other labels, such as selenomethionine for X-ray crystallography, selective labeling of different amino-acid positions using 1,3-^13^C-glycerol or 2-^13^C-glycerol or other precursors as carbon source, etc.

In summary, two unexpected and counterintuitive findings, the limited biomass generation and the irrelevance of the isotope integration step, have allowed the development of a highly optimized protocol that offers significant advantages over other published protocols: simplicity, no need for monitoring, minimal active time, high yields of labeled protein and cost reduction..

## Supporting information

Supplemental material. Protocol and Recipes

## Acknowledgements

The author thanks David Pantoja-Uceda for assistance with NMR spectroscopy and Douglas V. Laurents for constructive feedback. Proteins were expressed and NMR spectra recorded at the Manuel Rico NMR Laboratory (LMR), a node of the ICTS for biomolecular NMR (R-LRB).

## REFERENCES

Bok F, Moog HC, Brendler V (2023) The solubility of oxygen in water and saline solutions. Frontiers in Nuclear Engineering 2:. 10.3389/fnuen.2023.1158109

Cai M, Huang Y, Craigie R, Clore GM (2019) A simple protocol for expression of isotope-labeled proteins in Escherichia coli grown in shaker flasks at high cell density. J Biomol NMR 73:. 10.1007/s10858-019-00285-x

Kazmin D, Edwards RA, Turner RJ, et al (2002) Visualization of proteins in acrylamide gels using ultraviolet illumination. Anal Biochem 301:. 10.1006/abio.2001.5488

Lempp M, Farke N, Kuntz M, et al (2019) Systematic identification of metabolites controlling gene expression in E. coli. Nat Commun 10:. 10.1038/s41467-019-12474-1

Marley J, Lu M, Bracken C (2001) A method for efficient isotopic labeling of recombinant proteins. J Biomol NMR 20:. 10.1023/A:1011254402785

Marr AG (1991) Growth rate of Escherichia coli. Microbiol Rev 55:. 10.1128/mr.55.2.316-333.1991

Miller JH (1972) Experiments in molecular genetics. Cold Spring Harbor,NY: Cold Spring Harbor Laboratory

Neidhardt FC, Bloch PL, Smith DF (1974) Culture medium for enterobacteria. J Bacteriol 119:. 10.1128/jb.119.3.736-747.1974

Partridge JD, Sanguinetti G, Dibden DP, et al (2007) Transition of Escherichia coli from aerobic to micro-aerobic conditions involves fast and slow reacting regulatory components. Journal of Biological Chemistry 282:. 10.1074/jbc.M700728200

Sánchez-Clemente R, Igeño MI, Población AG, et al (2018) Study of pH Changes in Media during Bacterial Growth of Several Environmental Strains

Shiloach J, Fass R (2005) Growing E. coli to high cell density - A historical perspective on method development. Biotechnol Adv 23

Sivashanmugam A, Murray V, Cui C, et al (2009) Practical protocols for production of very high yields of recombinant proteins using Escherichia coli. Protein Science 18:. 10.1002/pro.102

