## Supplemental material. Protocol and Recipes for "Counter-intuitive method improves yields of isotopically-labeled proteins expressed in flask-cultured *Escherichia coli*"

**PROTOCOL**

"Unattended" Protocol for 13C-labelled protein expression.

- Day 1
  - grow pre-inoculum (around 1ml in LB) at 30-37ºC overnight. This step can be eliminated if enough cells from colonies in plate or glycerol frozen cells are directly used.
- Day 2.
  - Prepare modified M9++minimal medium (see recipe) and add **0.2% UNLABELED D-glucose** and 0.06% **LABELED** **^15^NH_4_Cl**.
  - Use an Tunair or Erlenmeyer flask with V≥ 10-20 V medium for the culture.
  - Inoculate with bacteria grown the day before to 0.05-0.1 OD_600_ (this is not too relevant).
  - Keep shaking vigorously (200rpm) at 25ºC for 24h.
- Day 3
  - OD_600_ should normally reach around 2.5 but it is no necessary to check.
  - Temper to the induction/expression temperature.
  - Add 1% **^13^C-D-glucose** and 0.3% **^15^NH4Cl** and IPTG according to your protocol.

Accustoming the cells to the minimal medium or waiting for isotope incorporation is unnecessary.

- o Maintain culture as usual (20ºC, 24h ensure complete consumption of ^13^C and its incorporation to the protein)
- Day 3 or 4 (depending on expression conditions).
  - Harvest cells as usual

**RECIPES**

100mL modified M9++ minimal medium (modified from Cai et al (2019)).

To 80ml H_2_O add:

- LB medium	100μL
- Trace elements^1^	20μL
- MgSO4 (1M)	100μL
- 100xBME vitamins^2^	250μL
- Thiamine (1mg/ml)	10μL
- CaCl2 (1M)	20μL
- 5xSalts M9++	20mL ALWAYS ADD AT THE END! (or at least after traces and CaCl2 to avoid precipitation) (salts recipe below)
- Antibiotic

Salts M9++ 5x

For 1L:

- K_2_HPO_4_	95g
- KH_2_PO_4_	25g
- Na_2_HPO_4_	45g
- K_2_SO_4_	12g

Trace elements solution (100mL):

- FeSO_4_ (7H2O)	0.6g
- MnCl_2_ (4H2O)	0.12g
- CoCl_2_ (6H2O)	0.08g
- ZnSO_4_ (7H2O)	0.07g
- CuCl2 (2H2O)	0.03g
- H_3_BO_4_	0.002g
- (NH_4_)_6_Mo_7_O_24_(4H2O)	0.025g
- EDTA	0.5g
